# The dual nature of metacommunity variability

**DOI:** 10.1101/2021.04.09.439168

**Authors:** Thomas Lamy, Nathan I. Wisnoski, Riley Andrade, Max C.N. Castorani, Aldo Compagnoni, Nina Lany, Luca Marazzi, Sydne Record, Christopher M. Swan, Jonathan D. Tonkin, Nicole Voelker, Shaopeng Wang, Phoebe L. Zarnetske, Eric R. Sokol

## Abstract

There is increasing interest in measuring ecological stability to understand how communities and ecosystems respond to broad-scale global changes. One of the most common approaches is to quantify the variation through time in community or ecosystem aggregate attributes (e.g., total biomass), referred to as aggregate variability. It is now widely recognized that aggregate variability represents only one aspect of communities and ecosystems, and compositional variability, the changes in the relative frequency of species in an assemblage, is equally important. Recent contributions have also begun to explore ecological stability at regional spatial scales, where interconnected local communities form metacommunities, a key concept in managing complex landscapes. However, the conceptual frameworks and measures of ecological stability in space have only focused on aggregate variability, leaving a conceptual gap. Here, we address this gap with a novel framework for quantifying the aggregate and compositional variability of communities and ecosystems through space and time. We demonstrate that the compositional variability of a metacommunity depends on the degree of spatial synchrony in compositional trajectories among local communities. We then provide a conceptual framework in which compositional variability of (i) the metacommunity through time and (ii) among local communities combine into four archetype scenarios: **spatial stasis** (low/low); **spatial synchrony** (high/low); **spatial asynchrony** (high/high) and **spatial compensation** (low/high). We illustrate this framework based on numerical examples and a case study of a macroalgal metacommunity in which low spatial synchrony reduced variability in aggregate biomass at the metacommunity scale, while masking high spatial synchrony in compositional trajectories among local communities. Finally, we discuss the role of dispersal, environmental heterogeneity, species interactions and suggest future avenues. We believe this framework will be helpful for considering both aspects of variability simultaneously which is important to better understand ecological stability in natural and complex landscapes in response to environmental changes.

## Introduction

Ecological stability is a fundamental concept to understand both current and future dynamics of ecosystems (MacArthur 1955, May 1973, Grimm and Wissel 1997, Ives and Carpenter 2007). While ecological stability may be quantified in various ways (Donohue et al. 2013, 2016, Hillebrand et al. 2018, Hillebrand and Kunze 2020, White et al. 2020), measures of variability through time are one of the most common approaches (Donohue et al. 2016, Xu et al. 2021). Temporal variability is usually measured by quantifying the temporal coefficient of variation of an aggregate attribute of an ecosystem, such as the total biomass of a given multispecies assemblage (hereafter referred to as aggregate community variability, see Table 1 for a glossary). Abundant empirical and theoretical evidence suggests that more taxonomically diverse communities exhibit lower aggregate variability (e.g., total biomass is less variable through time) due to the higher chance of a diverse community having species with redundant functional contributions to an ecosystem (i.e., compensatory dynamics) (Tilman 1999, Yachi and Loreau 1999, Gonzalez and Loreau 2009, Brown et al. 2016, Xu et al. 2021). However, communities do not exist in isolation; they are spatially connected via the dispersal of constituent species to form metacommunities over broader regional scales (Leibold et al. 2004, Leibold and Chase 2018). Such connectivity among local communities is important because it ultimately determines the temporal variability of the collection of communities at regional spatial scales (hereafter referred to as aggregate metacommunity variability, see Table 1; Wang and Loreau 2014, 2016). Understanding temporal variability over broader spatial scales at which metacommunities operate is key to managing complex landscapes, especially in the context of rapid environmental changes.

**Table 1.** Glossary

| Term | Definition |
| --- | --- |
| Local scale | Local communities delimited within “patches” (or sites) within the metacommunity. |
| Regional scale | The collection of all local communities. Also referred to as the metacommunity. |
| Metacommunity | A set of local communities connected by the dispersal of potentially interacting species. |
| Temporal variability | The fluctuation in time of a given attribute. Here, we focus on the temporal variability of both aggregate (e.g., total biomass) and compositional attributes. |
| Compositional variability | Variability in time in the relative frequencies of the species that make up local communities or the whole metacommunity. Independent of aggregate variability. |
| Aggregate variability | Temporal variability in the aggregate attribute of a community (i.e., total community biomass) or of the whole metacommunity (i.e., total metacommunity biomass). |
| Spatial *aggregate* synchrony | The degree of synchrony in an aggregate community attribute, usually total community biomass, among local communities. |
| Spatial *compositional* synchrony | The degree of synchrony in compositional change among local communities. |
| Environmental synchrony | The degree of synchrony in environmental change among local communities. |

Recent theoretical (Wang and Loreau 2014, 2016) and empirical (Wilcox et al. 2017, Wang et al. 2017, 2019) contributions have shown how considering aggregate variability from local to regional (i.e., the metacommunity) spatial scales is relevant for a richer understanding of ecological stability across space and time. Indeed, aggregate metacommunity variability (*i.e.*, temporal CV of total biomass summed across both species and local communities) critically depends on the degree of spatial “*aggregate*” synchrony among local communities (Table 1). For instance, if fluctuations in total biomass are spatially synchronous among local communities, aggregate metacommunity variability is also high (Figure 1A-B). Conversely, a low degree of spatial *aggregate* synchrony reduces aggregate metacommunity variability, despite potentially large aggregate variability at the local scale (*i.e.*, large CV of total biomass within each local community), and provides a spatial insurance effect (Loreau et al. 2003, Wang and Loreau 2014, 2016; Figure 1C-D). There have been few empirical examples of the mechanisms influencing metacommunity variability (e.g., Wilcox et al. 2017, Wang et al. 2019, 2021), partly due to a lack of theoretical development and long-term, broad-scale community datasets (Oliver et al. 2010, Donohue et al. 2013, Wang and Loreau 2014, 2016). The few empirical examples exploring such mechanisms have highlighted the importance of the taxonomic diversity among local communities (*i.e.*, beta diversity), which has the potential to reduce spatial *aggregate* synchrony and decrease temporal variability at the metacommunity scale. However, all of these prior studies solely focused on aggregate variability.

**Figure 1.**
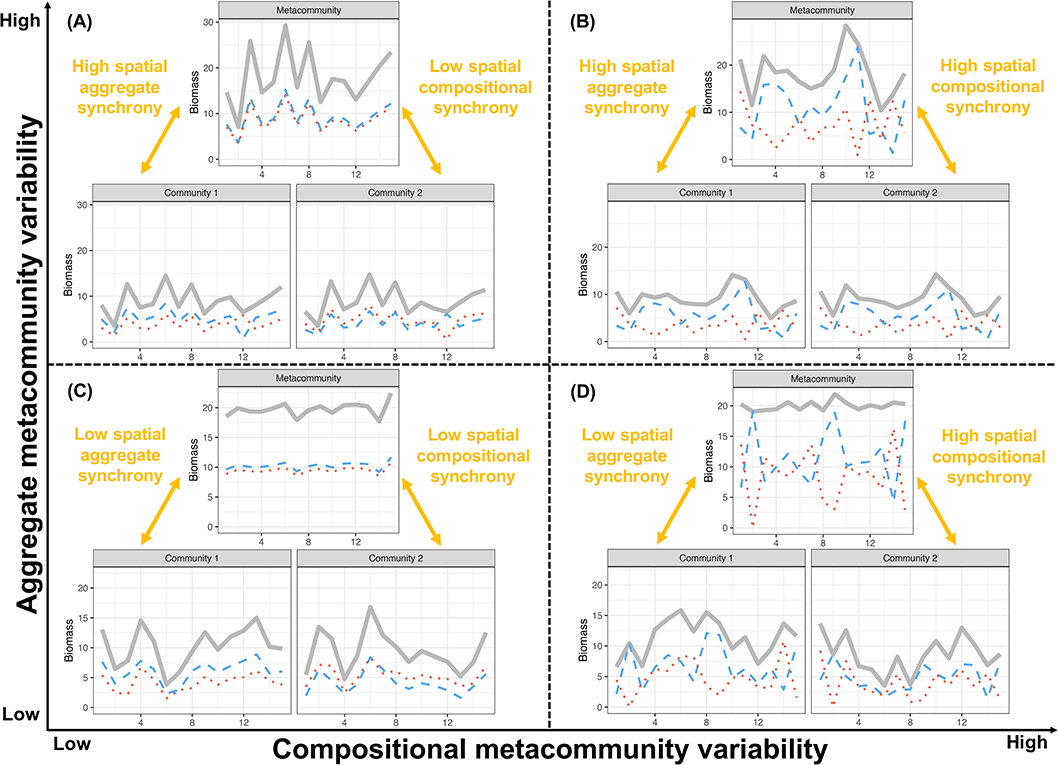
Conceptual illustration of the dual nature of metacommunity variability. Each panel displays a scenario of low (C, D) and high (A, B) aggregate metacommunity variability and low (A, C) and high (B, D) compositional metacommunity variability. Scenarios are based on two local communities (community 1 and 2) composed of two species surveyed for 15 years (x-axis of inset panels). Within each local community, dashed red and blue lines represent the biomass of the two species, and the solid grey line represents the total community biomass. At the metacommunity scale, dashed red and blue lines represent the metapopulation biomass of the two species, and the solid grey line the total metacommunity.

Aggregate variability represents only one facet of how communities and ecosystems can respond to environmental change, and *compositional* variability—change in the relative abundance or biomass of component species—is an equally important facet of variability (Micheli et al. 1999, Hillebrand et al. 2018, Hillebrand and Kunze 2020, White et al. 2020). The distinction between aggregate and compositional variability is important because compositional variability can beget or reduce aggregate variability. For instance, compositional variability reflecting rapid changes in species composition through time due to compensatory dynamics can directly decrease aggregate variability (Hillebrand et al. 2018). Although this dual nature of community variability is now well recognized and investigated in depth (e.g., White et al. 2020), the scaling of variability in space has only focused on aggregate variability leaving a conceptual gap in our understanding of temporal variability at the broader spatial scales at which metacommunities operate.

Building on these local-scale frameworks (Micheli et al. 1999, Hillebrand et al. 2018, Hillebrand and Kunze 2020), we address this knowledge gap by extending the concepts of aggregate and compositional variability to regional scales. This distinction is important because aggregate metacommunity variability can arise with or without compositional metacommunity variability. For instance, high aggregate metacommunity variability, as represented by large fluctuations in total biomass at the regional scale, may arise while the relative frequencies of constituent species remain constant (Fig. 1A) or change (Fig. 1B) through time. Similarly, low aggregate metacommunity variability can mask high compositional metacommunity variability. This masking occurs when the composition of the species that comprise the metacommunity changes over time, but these changing species assemblages continue to produce the same biomass. For instance, low spatial *aggregate* synchrony can stabilize total metacommunity biomass (*i.e.*, low aggregate metacommunity variability; Figure 1D). However, if one species becomes dominant over time at the metacommunity scale, this compositional change could remain undetected (Figure 1D), despite having important implications for the maintenance of biodiversity and species conservation at the regional scale (e.g., invasive species).

To provide a richer understanding of mechanisms underlying ecological stability through space and time, we provide a conceptual and methodological framework for quantifying aggregate and compositional variability through time at the local (*i.e.*, community) and regional (*i.e.*, metacommunity) scales. We propose a new way to partition compositional variability across spatial scales to compare the two facets of metacommunity variability. We then illustrate this framework based on numerical examples and a case study consisting of kelp forest communities in the Santa Barbara Channel off the coast of California, USA. We conclude with a conceptual framework for describing patterns of compositional metacommunity variability, in which compositional variability of (i) the metacommunity through time and (ii) among local communities combine into four archetype scenarios: (1) **spatial stasis**, low compositional metacommunity variability and low compositional variability among local communities; (2) **spatial synchrony**, high compositional metacommunity variability and low compositional variability among local communities; (3) **spatial asynchrony**, high compositional metacommunity variability and high compositional variability among local communities; (4) **spatial compensation**, low compositional metacommunity variability and high compositional variability among local communities. We discuss how dispersal, environmental heterogeneity, and species interactions can generate these scenarios and outline the general importance of better integrating this approach to understand ecological stability across spatial scales.

## Incorporating composition into temporal metacommunity variability

Variation in species composition has been studied extensively in a spatial context (Chase 2010, Anderson et al. 2011), and a common approach is to measure taxonomic beta diversity (Tuomisto 2010a, b, Anderson et al. 2011, Legendre and De Cáceres 2013). However, less attention has been given to variation in species composition through time (Adler et al. 2005, Hillebrand et al. 2010, Magurran et al. 2018, De Cáceres et al. 2019, Legendre 2019, Tatsumi et al. 2021).

### Summary of temporal beta diversity developments

Approaches to studying temporal taxonomic beta diversity have proliferated in recent decades. While descriptive and ordination-based approaches dominate the literature, new metrics are uncovering novel insights into community dynamics as multispecies time series increase in length and availability (Buckley et al. 2021). For example, studies have quantified the turnover in community composition between time points or relative to a baseline using dissimilarity metrics (Dornelas et al. 2014), shifts in species ranks in relative abundance (Avolio et al. 2019), and by partitioning compositional change into its turnover and nestedness components (Baselga 2010, Podani et al. 2013, Magurran et al. 2019).

Our lack of understanding of temporal taxonomic beta diversity at the regional scale of metacommunities presents an open challenge to identify the contributions of local and regional spatial processes to temporal variability (Magurran et al. 2019). The integration of spatial and temporal beta diversity has developed more slowly than spatial or temporal metrics alone, yet offers promise for understanding differences in temporal trajectories among communities (Legendre and Gauthier 2014), and how measures of synchrony influence regional dynamics (Hautier et al. 2018, Wang et al. 2021). Other approaches partition colonization and extinction dynamics that generate variation through time in spatially explicit landscapes (Tatsumi et al. 2020, 2021). While the best approach depends upon the question being asked, resolving how temporal beta diversity scales across space remains open for the development and examination of new and empirically testable approaches.

### Quantifying temporal compositional variability

Ideally, any metric of compositional variability should be independent of aggregate variability to reveal new insights not already captured by the latter and such metrics should be partitioned multiplicatively across spatial scales to allow for meaningful comparisons between aggregate and compositional variability. Here we build on the variance framework developed by Legendre and De Cáceres (2013). In this framework, beta diversity can be assessed based on any pairwise dissimilarity index (Anderson et al. 2011, Legendre and Legendre 2012), however, many of these indices depend on differences in aggregate attributes, such as total community biomass across samples (Legendre 2014, but see Lamy et al. 2015). To ensure that compositional variability is independent from aggregate variability our approach can only rely on pairwise dissimilarity indices based on species’ relative frequencies (Jost et al. 2011). Only three of these indices exist: Whittaker’s index of association (Whittaker 1952), the Chord distance (Orloci 1967) and the Hellinger distance (Rao 1995). For the partitioning of compositional variability, only the Hellinger and Chord distances are appropriate (Legendre and De Cáceres 2013). In the following, we develop an approach to partitioning compositional variability across space based on the Hellinger distance. The Hellinger distance is widely used in ecological studies (Legendre and Gallagher 2001) and closely related to the Chord distance — it is the Chord distance applied to square-root transformed species data — thus we based our approach only on Hellinger distance for the sake of clarity.

The compositional variability of a single community *i* composed of *s* species surveyed *n* times can be computed as beta diversity (BD) based on the variance framework of Legendre and De Cáceres (2013). In this framework, the compositional variability of community *i* is the total variance of species composition through time computed as:

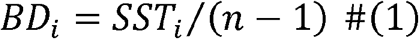

where *SST_i_* is the total sum of squares in species composition, 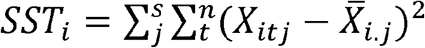, with 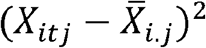 the square of the difference between the biomass of species *j* at time *t*, and the temporal mean biomass of species 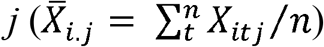. This definition corresponds to the Euclidean distance, which is inappropriate to assess beta diversity (Wolda 1981, Legendre and Gallagher 2001) and does not fulfill the density invariance property described previously (Jost et al. 2011). For appropriate calculation of compositional variability and meaningful comparisons with aggregate variability, we suggest computing *SST* corresponding to the Hellinger distance.

Compositional variability based on the Hellinger distance (*BD^h^*; where *h* stands from Hellinger) is calculated by applying equation 1 to the square root of species relative frequencies (*i.e.* the Hellinger transformation of the original data). Compositional variability of community *i* based on the Hellinger distance can be rewritten as:

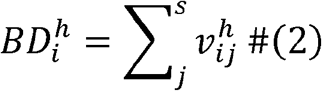

where 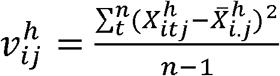 is the temporal variance of the Hellinger-transformed biomass of species *j* in the local community *i*. Here,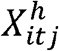 represents the Hellinger transformation of the biomass of species *j* in community *i* at time *t* (Table 2). Greater sums of species variances 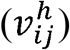 lead to greater compositional variability 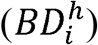.

**Table 2.** Notations summary for aggregate and compositional metacommunity variability across spatial scales. Note that “.” are used to define sums (e.g., 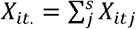)

| Symbol | Description |
| --- | --- |
| $X_{itj}$ | Biomass of species $j$ in community $i$ at time $t$ |
| $X_{it.} = \sum_j^s X_{itj}$ | Total biomass of community $i$ at time $t$ |
| $X_{itj}^h = \sqrt{X_{itj}/X_{it.}}$ | Hellinger transformation of the biomass of species $j$ in community $i$ at time $t$, corresponding to the square root coefficient of species relative frequency |
| $X_{i..} = \sum_t^n \sum_j^s X_{itj}$ | Total biomass of community $i$ |
| **Local scale aggregate variability** |  |
| $\sigma_{Ti}$ | Temporal standard deviation of total biomass in community $i$ |
| $CV_\alpha^2 = \left( \frac{\sum_i^m \sigma_{Ti}}{\mu_{TT}} \right)^2$ | Local scale aggregate variability, defined as the square coefficient of the weighted average of aggregate variability across communities |
| **Local scale compositional variability** |  |
| $v_{ij}^h = \frac{\sum_t^n (\sqrt{X_{itj}/X_{it.}} - \sqrt{\overline{X_{itj}/X_{it.}}})^2}{n-1}$ | Temporal variance of the Hellinger-transformed biomass of species $j$ in community $i$ |
| $BD_i^h = \sum_j^s v_{ij}^h$ | Compositional variability of community $i$, corresponding to the beta diversity based on the Hellinger distance of community $i$ |
| $BD_\alpha^h = \sum_i^m \frac{X_{i..}}{X_{...}} \sum_j^s v_{ij}^h$ | Local scale compositional variability, defined as the weighted average of compositional variability across communities |
| **Regional scale aggregate variability** |  |
| $\sigma_{TT}$ | Temporal standard deviation of the total biomass of the whole metacommunity |
| $\mu_{TT}$ | Temporal mean of the total biomass of the whole metacommunity |
| $CV_\gamma^2 = \left( \frac{\sigma_{TT}}{\mu_{TT}} \right)^2$ | Metacommunity aggregate variability |
| **Regional scale compositional variability** |  |
| $X_{.tj} = \sum_i^m X_{itj}$ | Metacommunity biomass of species $j$ at time $t$ |
| $X_{.t.} = \sum_i^m \sum_j^s X_{itj}$ | Total metacommunity biomass at time $t$ |
| $v_{Tj}^h = \frac{\sum_t^n (\sqrt{X_{.tj}/X_{.t.}} - \sqrt{\overline{X_{.tj}/X_{.t.}}})^2}{n-1}$ | temporal variance of the Hellinger-transformed metacommunity biomass of species $j$ |
| $BD_\gamma^h = \sum_j^s v_{Tj}^h$ | Metacommunity compositional variability corresponding to the beta diversity based on the Hellinger distance of the whole metacommunity |
| **Spatial aggregate and compositional synchrony** |  |
| $\varphi = \left( \frac{\sigma_{TT}}{\sum_i^m \sigma_{Ti}} \right)^2$ | Spatial aggregate synchrony defined following Wang and Loreau (2014) and Loreau and de Mazancourt (2008) |
| $BD_\varphi^h = \frac{\sum_j^s v_{Tj}^h}{\sum_i^m \frac{X_{i..}}{X_{...}} \sum_j^s v_{ij}^h}$ | Spatial component scaling compositional variability between $BD_\alpha^h$ and $BD_\gamma^T$. Does not correspond to the synchrony definition of Loreau and de Mazancourt (2008) |

## Linking aggregate and compositional variability across spatial scales

Both aggregate and compositional metacommunity variability (γ variability) can be multiplicatively partitioned into local-scale variability (α) and a spatial component (φ). The spatial component (φ) corresponds to the spatial *aggregate* synchrony and the spatial *compositional* synchrony that quantify how aggregate and compositional variability, respectively, scale up from the local scale to the whole metacommunity (Table 1). Aggregate metacommunity variability 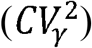 can be multiplicatively partitioned as 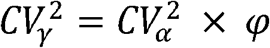 (Wang and Loreau 2014). 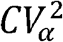 is the average aggregate community variability at the local scale and φ is the spatial *aggregate* synchrony (Table 1 and 2). φ ranges between zero and one, with higher φ indicating that fluctuations in total biomass are spatially synchronous among local communities. Here we present a similar approach to partitioning compositional metacommunity variability based on the Hellinger distance.

For a given metacommunity, we assume each local community is sampled in a similar way, such that *m* communities (or sites) are sampled over *n* time steps. During each survey, the biomass of *s* species is recorded. The data can be summarized as a community array **X**, where *X_itj_* represents the biomass of species *j* in community *i* at time *t*. The metacommunity corresponds to the broadest spatial scale and is defined by (i) the total metacommunity biomass obtained by summing the biomass across all communities and species and (ii) a *n* x *s* time-by-species matrix containing the regional biomass of each species over time obtained by summing the biomass across all communities (Table 2).

### Local scale compositional variability

Mean local scale compositional variability is computed as the weighted average of compositional variability across the *m* communities calculated using 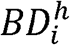

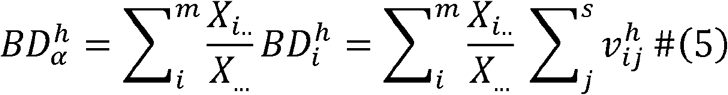

with 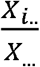 the weight of local community *i*. Local communities are weighted by their contributions to the overall metacommunity biomass with 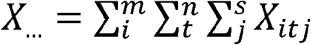and 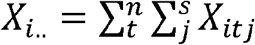 In eq. 5, 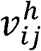 represents the temporal variance of the Hellinger-transformed biomass of species *j* in the local community *i*.

### Regional scale compositional variability

We define the regional scale compositional variability as:

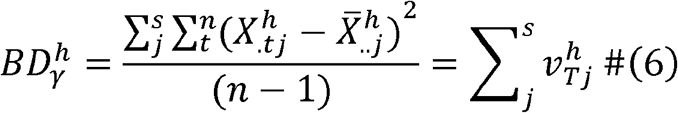

where 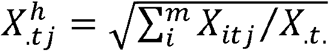 is the square root of the regional frequency of species *j* at time *t*, and 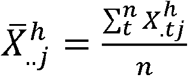 is the temporal mean of the square root coefficient of the regional frequencies of species 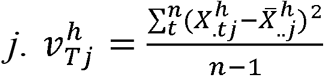 corresponds to the temporal variance of the Hellinger-transformed regional biomass of species *j*. Thus, greater variability of individual species frequencies at the regional scale contributes to greater regional scale compositional variability (i.e., larger values of 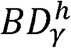).

### Linking compositional variability across multiple spatial scales: the spatial synchrony components

Similar to aggregate variability, we propose that compositional variability at the regional scale 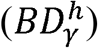 can be partitioned multiplicatively into a local scale 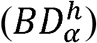 and a spatial component 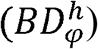 as 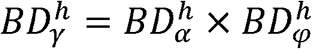. Spatial *compositional* synchrony, 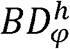, reflects how compositional variability scales from local communities to the metacommunity. 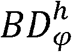 increases as compositional trajectories among communities become more spatially synchronous. The spatial component 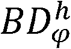 is defined as the ratio between gamma and alpha compositional variability:

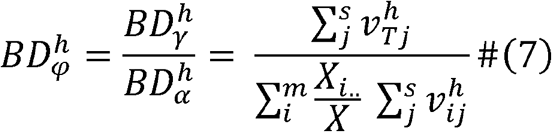

The quantitative partitioning of variability into local community-scale (α), regional metacommunity-scale (γ), and spatial (φ) components of aggregate and compositional variability suggests the existence of a common currency to investigate metacommunity variability. Notably, φ and 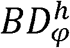 are essential components to understand ecological stability across spatial scales.

These metrics can be calculated using the *metacommunity_variability* function in the *ltmc* (long-term metacommunity analysis) package for R (available at https://github.com/sokole/ltermetacommunities/tree/master/ltmc). See supplemental material SM1 for a worked example.

## Illustrations of compositional and aggregate variability across space and time

### Simulated examples

Here we present simulated examples to further understand and illustrate what φ and 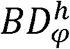. The first example encompasses the four scenarios presented in Figure 1 for two local communities composed of two species each surveyed for 15 years. Each scenario was built by simulating correlated species biomasses (dashed red and blue lines) within communities and correlated total biomasses (solid grey lines) among communities based on the Cholesky factorization method. In Figure 1A, species biomasses were positively correlated (ρ = 0.95) and mirrored across the two communities (species biomasses across two mirrored communities change in such a way that species relative frequencies at the regional scale is constant through time). Total biomass was positively correlated (ρ = 0.95). Intuitively, high aggregate metacommunity variability 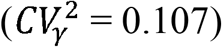 was explained by high spatial *aggregate* synchrony among communities (φ = 0.981), whereas low compositional metacommunity variability 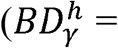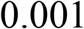; i.e., low variability in species relative frequencies at the regional scale) was explained by low spatial *compositional* synchrony among communities 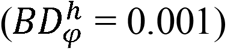. In Figure 1B, species biomasses were negatively correlated (ρ = 0.95) but identical across communities. Total biomass was positively correlated (ρ = 0.95). Both aggregate 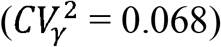 and compositional 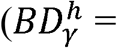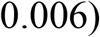 metacommunity variability were high due to large degree of both spatial *aggregate* (φ = 0.965) and *compositional* 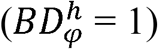 synchrony. In this case, variability at the regional (*i.e.*, metacommunity) scale mimics variability at the local (*i.e.*, community) scale (i.e., 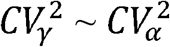 and 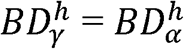). In Figure 1C, species biomasses were positively correlated with one another within patches (ρ = 0.95) and mirrored across communities, while total biomass was negatively correlated between communities (ρ = −0.95). Both aggregate 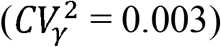 and compositional 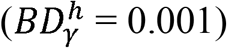 metacommunity variability were low due to a small degree of spatial *aggregate* (φ = 0.0031) and *compositional* 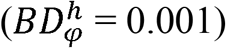 synchrony. In Figure 1D, species biomasses were negatively correlated (ρ = −0.95) and identical across communities, while total biomass was negatively correlated (ρ = −0.95). Low aggregate metacommunity variability 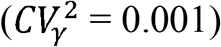 was explained by a low amount of spatial *aggregate* synchrony (φ = 0.016) while high compositional metacommunity variability 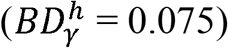 was explained by a higher degree of *compositional* synchrony (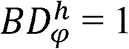 and therefore 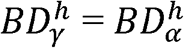).

We ran additional simulations to illustrate which aspect of compositional variability was captured by 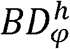 and test whether 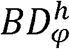 was independent from fluctuations in aggregate attributes (e.g., total biomass). The next three examples consist of two local communities (community 1 and 2) composed of four species surveyed for 15 years (Figure 2A). Species relative frequencies at the metacommunity scale were constant across species and over time (p = 0.25). We then randomly generated species relative frequencies in community 1 and inferred those in community 2 as the difference between community 1 and the metacommunity (p = 0.25). Consequently, the compositional trajectory of community 2 mirrors that of community 1, so that their species relative frequencies are spatially anti-correlated over time and there are no compositional changes through time at the metacommunity scale. We computed φ and 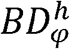 for a metacommunity consisting of only communities 1 and 2, and iteratively increased the size of the metacommunity by adding a community identical to community 2 over 100 iterations. Thus, with each additional community mimicking community 2, the metacommunity should become more spatially synchronous in composition, which 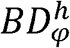 should detect. The total biomass of each community was either constant (Figure 2A) or randomly chosen (Figure 2B) at each iteration. Intuitively, 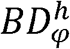 increased as the number of the local communities exhibiting similar compositional trajectories increases in the metacommunity, before plateauing to ∼0.95 (Figures 2A and 2B). When the total biomass in communities 1 and 2 remained identical across iterations (Figure 2A), φ also increased asymptotically as a result of adding communities with similar fluctuations in total biomass. However, when the total biomass in communities 1 and 2 were randomly drawn at each iteration φ quickly dropped to zero (Figure 2B) since the random draws generated independent fluctuations in total biomass as communities were added to the metacommunity. Our final example consists of a metacommunity with two communities with mirrored compositional trajectories, but with a standard deviation σ*_TT_* (Table 2) in total metacommunity biomass increasing by 0.25 at each iteration (from 1 to 30). In this scenario, we would want to separately detect the low compositional variability (shown by the constant species relative frequencies at the regional scale) from increasingly large variability in aggregate attribute. This example shows that 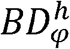 was not influenced by increasing aggregate variability and remained near zero as the standard deviation in total metacommunity biomass increased (Figure 3C), whereas φ increased.

**Figure 2.**
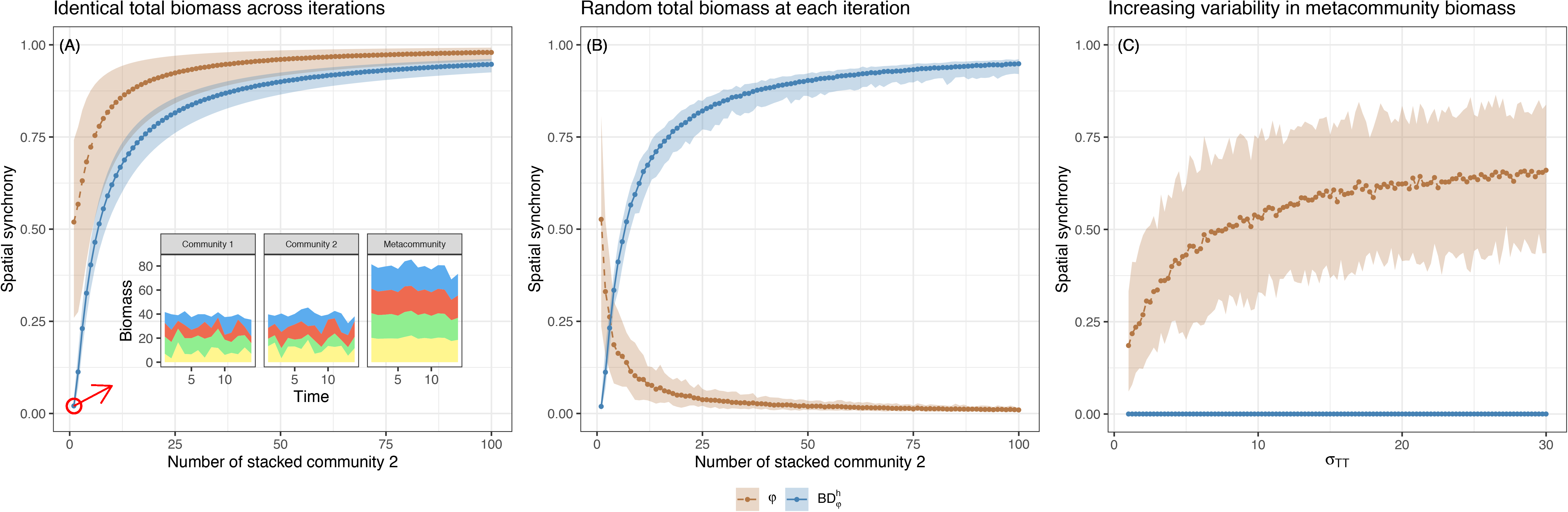
Numerical examples consisting of two local communities (community 1 and 2) composed of four species surveyed for 15 years. In each case species relative frequencies in community 1 and 2 were randomly generated so their compositional trajectories are mirrored. Therefore, compositional variability at the metacommunity scale was null (species relative frequency at the metacommunity scale was 0.25 for each species at each time step; see inset of A for an example). (A) A metacommunity of increasing size generated by stacking community 2 over 100 iterations. The total biomass of community 1 and 2 was constant across iterations. (B) A metacommunity of increasing size generated by stacking community 2 over 100 iterations. The total biomass of each community was randomly chosen at each iteration. (C) A metacommunity with only two mirrored communities 1 and 2 and a standard deviation σ*_TT_* in total metacommunity biomass increasing by 0.25 at each iteration (from 1 to 30). All scenarios were generated 99 times and mean φ and 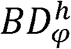 along with their 95% confidence intervals were reported.

**Figure 3.**
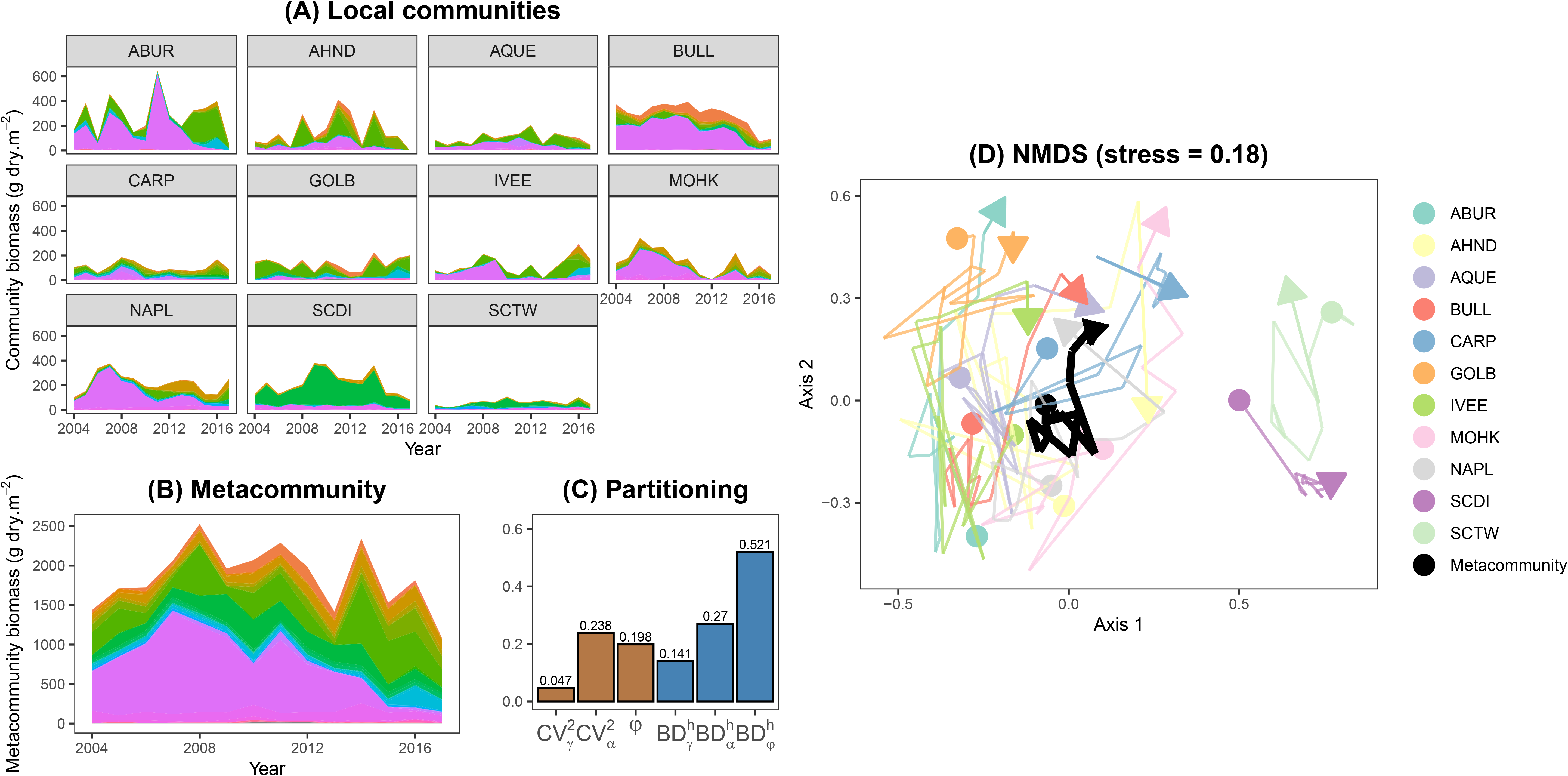
Case study of macroalgae that inhabit shallow rocky reefs (data package ID: knb-lter-sbc.50.7). Representation of the community structure of (A) the 11 rocky reefs and (B) the metacommunity. Each color corresponds to one species. (C) Results from the partitioning of aggregate and compositional variability. Each color represents one method. (D) Compositional trajectories of rocky reef and the metacommunity based on NMDS. The compositional trajectory of the metacommunity is pictured as a thick black line and each color represents the compositional trajectory of one of the 11 rocky reefs.

These simulations illustrate that φ and 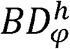 can capture patterns of spatial correlation in the aggregate (i.e., total biomass) and in the compositional trajectories, respectively, across a set of local communities. Importantly, φ and 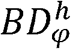 displayed independent patterns suggesting that these metrics capture different aspects of metacommunity variability.

### Case study: understory macroalgal communities

To illustrate the aggregate and compositional variability across space and time with empirical data, we focused on understory macroalgal communities inhabiting shallow rocky reefs off the coast of Santa Barbara, California, USA. From late July to early August of each year, the abundance of 55 macroalgae were recorded at 36 fixed 80 m^2^ plots distributed across 11 shallow (4 – 12 m depth) rocky reefs. Abundances were converted to biomass density (g decalcified dry mass m^−2^) using species-specific allometries (Harrer et al. 2013, Reed 2018). Data were collected annually from 2004 to 2017 as part of the Santa Barbara Coastal Long Term Ecological Research program (http://sbc.lternet.edu; Reed 2018) and are publicly available on the EDI Data Portal (https://doi.org/10.6073/pasta/d5fd133eb2fd5bea885577caaf433b30). We averaged species biomasses across the 11 rocky reefs (Figure 3A) and computed both aggregate and compositional variability of the metacommunity, which was obtained by summing species biomasses across the 11 reefs (Figure B). We then multiplicatively partitioned both aggregate and compositional metacommunity variability using the methods described above (Figure 3C).

We found that aggregate variability (i.e., fluctuations in total biomass) was reduced by a factor of ∼5 from the local (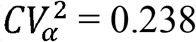; Figure 3A & 3C) to the metacommunity scale (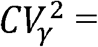 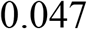; Figure 3B & 3C). This reduction occurred due to a relatively small degree of spatial *aggregate* synchrony (φ = 0.198), suggesting that the fluctuations in total biomass were weakly correlated across the 11 rocky reefs. However, compositional variability only decreased by a factor of ∼2 from the local scale 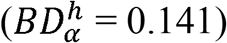 to the whole metacommunity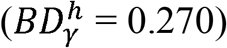 due to a higher degree of spatial *compositional* synchrony (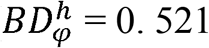; Figure 3C). Local communities in each rocky reef generally followed similar trends in composition over time (Figure 3A), which translated into higher compositional variability at the metacommunity scale (Figure 3B).

To further investigate compositional variability, we performed a nonmetric multidimensional scaling (NMDS) of Hellinger distances to assess the compositional trajectories of the 11 rocky reefs and of the metacommunity simultaneously over the 14-year period (Figure 3D). The NMDS provides evidence for both substantial compositional differences among rocky reefs (i.e., spatial taxonomic beta diversity) and temporal changes within each of these reefs. Many rocky reefs had relatively similar composition at the beginning of the survey (in the lower left quadrant of the NMDS plot). Over the 14-year monitoring period, most reefs experienced comparable compositional trajectories along the second axis (from the lower to the upper section of the plot; Figure 3D). As a result, the overall metacommunity trajectory tracked these local compositional changes.

## Compositional insight into metacommunity variability

Scaling variability from local to regional scales poses an exciting challenge for ecologists. Recent contributions (Wang and Loreau 2014, 2016, Wang et al. 2019) have provided the theoretical foundation for empirical investigations of aggregate variability across space and time (Wilcox et al. 2017, Wang et al. 2019, 2021). Yet, focusing only on aggregate variability overlooks a key component of metacommunity dynamics—compositional variability (Micheli et al. 1999, Hillebrand et al. 2018, Hillebrand and Kunze 2020). The framework presented here helps fill this knowledge gap and provides a first approach to quantify both the aggregate and compositional facets of metacommunity variability. In particular, our framework can lead to four extreme patterns of metacommunity variability that arise depending on the high or low values of aggregate metacommunity variability and compositional metacommunity variability introduced in Figure 1. These four scenarios are the direct translations of those found in Micheli et al. (1999), but at the metacommunity scale rather than the local community scale.

However, our framework differs from that of Micheli et al. (1999) in that metacommunity dynamics cannot be understood without an explicit consideration of the spatial dynamics across local communities. For instance, low compositional metacommunity variability can arise due to either (*i*) similarly low compositional variability within each local community or (*ii*) high compositional variability that are weakly synchronous across local communities (Figure 1A). Therefore, to understand the implications of considering compositional variability, we suggest to place metacommunities in a two-dimensional space defined by two properties: the compositional variability of the metacommunity thought time and the compositional variability among local communities (Figure 4). The second property actually tracks if compositional differences among communities remain the same over time (low), increase (high) or decrease (high) over time and allows to distinguish between cases (*i*) and (*ii*). We identify four archetype scenarios at the extreme ends of these continuums: (1) **spatial stasis**, low compositional metacommunity variability and low compositional variability among local communities through time; (2) **spatial synchrony**, high compositional metacommunity variability and low compositional variability among local communities through time; (3) **spatial asynchrony**, high compositional metacommunity variability and high compositional variability among local communities through time; (4) **spatial compensation**, low compositional metacommunity variability and high compositional variability among local communities through time.

**Figure 4.**
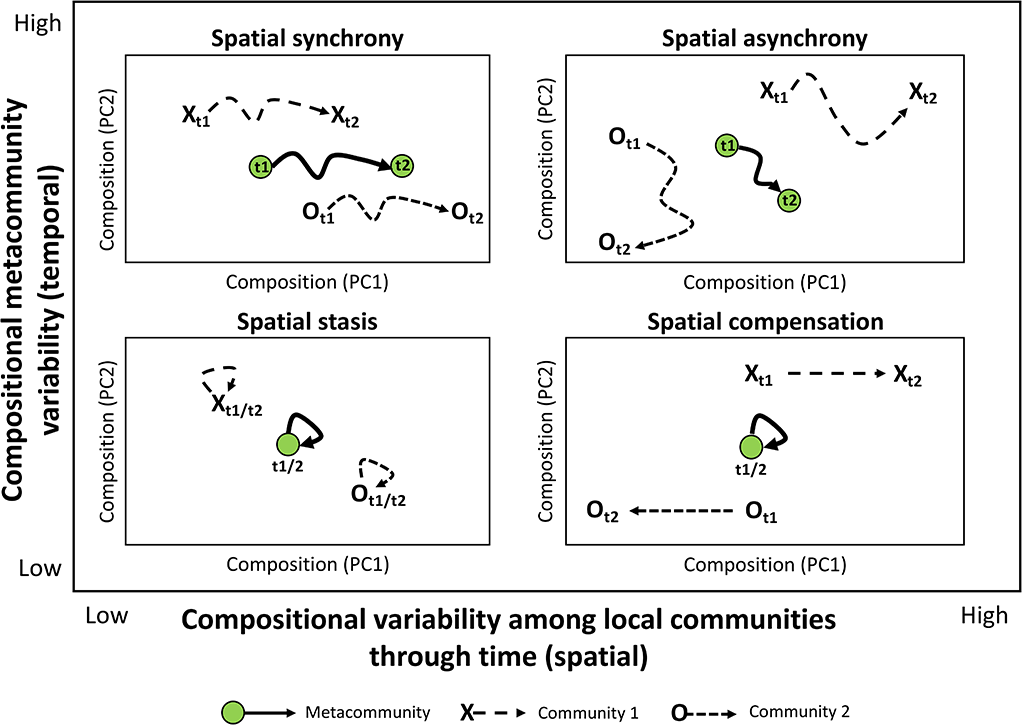
Conceptual framework for understanding the implications of considering compositional variability. Each sub-figure represents the temporal trajectory over nine years (from t1 to t9) of a metacommunity composed of two local communities (X and O) in a two-dimensional compositional space. We identified four archetype scenarios at the extreme ends of a two-dimensional space defined by two properties: the compositional variability of the metacommunity thought time (y-axis) and the compositional variability among local communities (x-axis) The four scenarios correspond to: (1) **spatial stasis**, low compositional metacommunity variability and low compositional variability among local communities through time; (2) **spatial synchrony**, high compositional metacommunity variability and low compositional variability among local communities through time; (3) **spatial asynchrony**, high compositional metacommunity variability and high compositional variability among local communities through time; (4) **spatial compensation**, low compositional metacommunity variability and high compositional variability among local communities through time.

In our empirical study of an understory marine macroalgal metacommunity (Figure 3) we found that compositional variability of the metacommunity was relatively high and exhibited large shifts in species assemblages that would have been undetected from investigating total biomass alone (Figure 3B). This scenario is akin to our scenario in Figure 1D and demonstrates how lack of variability in aggregate metacommunity attributes can mask compositional changes across space and time. Second, we found relatively high degree of *compositional* spatial synchrony. This result shows that most local communities underwent similar compositional trajectories, in particular, major declines of the dominant species that were partially compensated for by the increase of the three sub-dominant species (Lamy et al. 2019). Our empirical case study is therefore akin to the **spatial synchrony** scenario displayed in Figure 4. Further, our case study also illustrates how compositional variability contributes to the lack of aggregate variability at the metacommunity scale. Overall, the case study shows how understanding synchrony in the compositional dynamics across local communities may enable greater insights into the mechanisms underlying metacommunity variability.

## Mechanisms underlying metacommunity variability

As more long-term spatio-temporal surveys become available (Hughes et al. 2017, Record et al. 2021), it will become increasingly feasible to gain new insights into the mechanims underlying metacommunity variability across systems. Nonetheless, we can assume the four scenarios presented in Figure 4 result from a variety of ecological processes at play both within and among local communities. Notably, spatial *compositional* synchrony is directly influenced by the combined effects of three mechanisms: species interactions, dispersal, and environmental variation in space and time in the metacommunity (Amarasekare 2003, Leibold et al. 2004, Shoemaker and Melbourne 2016).

### Mechanisms underlying compositional metacommunity variability

**Spatial stasis** occurs when the metacommunity is characterized by relatively stable local environmental conditions that stabilize local species composition. This provides a baseline in the absence of disturbance and environmental change, and likely presents a scenario when populations of species within the local communities are at demographic equilibrium. Spatial stasis may also be more common for metacommunities characterized by long-lived organisms that are slow to change in composition. Spatial stasis is likely to become increasingly uncommon with global climate change and intensifying anthropogenic activities.

**Spatial synchrony** can occur when dispersal rates are high, leading to similar compositional trajectories across local communities, which ultimately increases compositional metacommunity variability (Gouhier et al. 2010). Alternatively, region-wide environmental forcing can also induce spatial synchrony in community dynamics (Steiner et al. 2013). For example, if changes in local environmental conditions are similar across patches, local community composition might follow similar trajectories if community assembly is influenced more by environmental drivers than biotic interactions, leading to high compositional metacommunity variability. This is probably the mechanism driving the metacommunity change observed in the case study of understory algae, as the consistent replacement of the dominant species by a few sub-dominant species was linked to broad-scale variations in temperature and nutrients along the California coast (Lamy et al. 2019). Spatial synchrony can also result from the regional synchronizing effect of highly mobile consumers, although this mechanism has been less well-described than dispersal and synchronizing environmental fluctuations (Ims and Steen 1990; de Roos et al. 1998). **Spatial asynchrony** can occur if environmental change is not strongly correlated in space, if dispersal is limited, and local biotic interactions strongly shape community assembly, or if community dynamics are largely stochastic. In this case, spatial *compositional* synchrony may be low, resulting in reduced compositional metacommunity variability. **Spatial compensation** represents an extreme case of spatial asynchrony in which the combination of different environmental conditions across local communities and limited dispersal shift local community composition in divergent directions. Thus, metacommunity composition may be stabilized by low-to-intermediate dispersal rates, low environmental variability over time, or a lack of spatial synchrony in environmental variability that reduces spatial *compositional* synchrony (Chalcraft 2013).

### Mechanisms underlying distinct aggregate and compositional responses

#### Low compositional metacommunity variability

If aggregate metacommunity variability is low, then the metacommunity can be considered stable and investigation into synchrony patterns could help decipher the actual mechanism at play. However, if aggregate metacommunity variability is high we can suppose that some aspect of the environment (e.g., disturbance, climatic change) limits function (i.e., biomass production) uniformly across all species through time. In both cases, lack of regional changes in species composition can occur either due to **spatial stasis** or **spatial compensation** (Figure 4) and discriminating between these two scenarios would require investigating the compositional dynamics across local communities.

#### High compositional metacommunity variability

This case is not necessarily destabilizing as compositional variability can contribute to low aggregate variability (e.g., in standing biomass) at broader spatial extents as outlined by our case study (Figure 3). In the case of **spatial synchrony**, similar compositional trajectories across communities can ensure ecosystem function (e.g., biomass production) remains high and stable at broader spatial extents where metacommunities operate. This, however, assumes that local compositional trajectories are due to compensatory dynamics (*sensu* Micheli et al. 1999). If stochasticity dominates, competitive exclusion by productive species is strong, or environmental changes favor less productive species, then resulting compositional changes within local communities can result in variable biomass production at the regional scale (if spatial synchrony is high).

### Synchrony in environmental change

Spatially autocorrelated environmental changes may increase metacommunity variability by increasing spatial aggregate and compositional synchrony. Any disturbance that increases the synchrony in environmental fluctuations will destabilize communities at broad spatial extents (Moran 1953). Therefore, the spatial scale of shared environmental fluctuations determines how many patches of the metacommunity are likely to be experiencing similar environmental conditions at any given time. This suggests a distinction between local-extent fluctuations and regional-extent fluctuations, such that regional-extent fluctuations may increase metacommunity variability more strongly than fluctuations at the local scale by inducing spatial synchrony in a larger portion of the metacommunity (Ruhi et al. 2018). Considering that it is much easier to measure environmental variation than species interactions and dispersal, it is generally easy to link aggregate and compositional responses to the degree of spatial autocorrelation in environmental variation. Indeed, sets of environmental variables can now be easily retrieved at local and regional scales. Satellite data are more than ever available to a broad audience (Nguyen et al. 2018, Bell et al. 2020) and environmental data layers are easily accessible through open-access datasets such as Bio-Oracle in the marine realms (Assis et al. 2018). Moreover, geographically extensive time series data, such as those gathered by long-term ecological research programs, have recently reached the multi-decadal durations suitable for making inferences about metacommunity synchrony through time (Clutton-Brock and Sheldon 2010, Edwards et al. 2010). We advocate for widespread integration of synchrony in environmental change to our proposed framework. This is particularly important because recent studies suggest that some systems are becoming more synchronous due to climate trends (Post and Forchhammer 2004), which may synchronize variability among communities and destabilize metacommunities.

## Implications and future avenues

### Implication for conservation

Our framework enables consideration of both aggregate and compositional variability across space and time and is thus relevant for conservation planners who are increasingly tasked with implementing local- and regional-scale strategies to minimize biodiversity loss (e.g., Gimona et al. 2012, Socolar et al. 2016). While the importance of spatial processes (e.g., colonization–extinction and source–sink dynamics) for conservation planning has been widely acknowledged (Margules and Pressey 2000), explicit use of the metacommunity concept by conservation practitioners has been rare, but effective. For example, application of the metacommunity framework to lowland heathland conservation showed that coordinated efforts among local sites could increase regional-scale conservation success (Diaz et al. 2013). In addition, this study recognized the importance of suboptimal patches for the maintenance of regional biodiversity, an aspect of conservation that is often overlooked, but would be important for identifying which and how many sites should be targeted for conservation (Socolar et al. 2016).

### Methodological considerations

Our main goal was to provide the conceptual foundation for the integration of an important, yet overlooked, facet of metacommunity variability: composition. Through this exercise we also provided the first empirical way to assess compositional variability across spatial scales based on the variance framework of Legendre and De Cáceres (2013).

A limitation of this approach is that the spatial *compositional* synchrony index 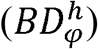 is only defined as the ratio between 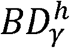 and 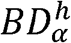 and can therefore exceed one. This is due to the fact that unlike aggregate variability (Wang and Loreau 2014) (or other metrics such as species richness), compositional variability defined as 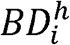 does not necessarily decrease with increasing spatial extent and therefore edge cases exist where _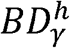_ can be smaller than 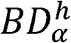. When shifting from a single local community to a collection of them (i.e., the metacommunity), species relative frequencies does not always increase as total biomass does. For instance, if there are two local communities made of two distinct species fluctuating over time, then 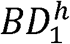 and 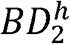of communities 1 and 2, respectively, will both be null (since there is a single species in each community its relative frequency is always 1), but 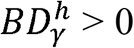. This is of course an extreme case, and our simple numerical examples suggest 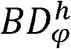 can adequately capture synchronous compositional trajectories among local communities. More generally it could be that 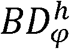 is greater than one when different sets of local communities have no taxa in common, indicating dissimilarity saturation (Tuomisto et al. 2012), but further work will be needed to ascertain this claim.

Assessing how compositional variability changes across space and time is a new and challenging topic and much work remains to be done. First, alternative metrics of spatial *compositional* synchrony should be investigated. Recent developments have relied on comparative geometry of community trajectories in multivariate space to quantify the convergence, divergence, or cyclic nature of temporal changes (De Cáceres et al. 2019, Sturbois et al. 2021) and represent a great avenue. Second, given that various issues have complicated the field of compositional variability in space (Jost 2007, Tuomisto 2010a, b), further simulations beyond those presented in this paper will be needed to assess these issues for different approaches that investigate compositional variability across time and space. Finally, the combination of 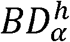, 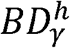 and 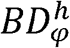 is just one synthetic way to investigate compositional variability across spatial scale. Other multivariate statistical methods should be used in complement to further scrutinize this facet (Legendre and Gauthier 2014, Lamy et al. 2015).

## Conclusions

Metacommunity variability through time has two complementary dimensions: aggregate and compositional. Wang et al. (2019) provided an integrative framework in which aggregate metacommunity variability can be partitioned either (*i*) from individual local populations to local communities and from local communities to the metacommunity or (*ii*) from individual local populations to metapopulations and from metapopulations to the metacommunity. The metric of spatial *compositional* synchrony we presented here, 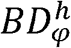, is a more integrative measure that directly quantifies how variability scales from local-scale populations to the metacommunity. It aims to capture the degree of synchrony in the compositional trajectories among local communities, thus providing complementary insights into the mechanisms that decrease temporal variability over broad spatial extents.

Partitioning aggregate and compositional variability across spatial scales yields a quantitative estimate of the degree of spatial *aggregate* synchrony and spatial *compositional* synchrony, thus facilitating the comparison between the two dimensions of community variability across spatial scales. Our framework links aggregate variability, based on previous work, and compositional variability across spatial scales to deepen our understanding of the mechanisms underlying metacommunity variability. The marine metacommunity case study illustrates how the joint focus on aggregate and compositional variabilities reveals important compositional changes at broad spatial extents, mainly due to synchronous compositional trajectories among local communities. This insight would have been overlooked if the focus had been solely on the aggregate components of the metacommunity. Our approach contributes to relevant conservation and management issues by yielding insight into the ecological processes that may stabilize or destabilize aspects of biodiversity at broad spatial extents.

## Supplementary material

SM1 (Supplementary material 1): R code to reproduce the example in Figure 3 is available at https://github.com/sokole/ltermetacommunities/tree/master/Manuscripts/MS2-Supp-Info/supp-info-example-agg-by-site-rev2019112

## Literature cited

Adler, P. B. et al. 2005. Evidence for a general species–time–area relationship. - Ecology 86: 2032–2039.

Amarasekare, P. 2003. Competitive coexistence in spatially structured environments: a synthesis. - Ecol. Lett. 6: 1109–1122.

Anderson, M. J. et al. 2011. Navigating the multiple meanings of β diversity: a roadmap for the practicing ecologist. - Ecol. Lett. 14: 19–28.

Assis, J. et al. 2018. Bio-ORACLE v2.0: Extending marine data layers for bioclimatic modelling. - Glob. Ecol. Biogeogr. 27: 277–284.

Avolio, M. L. et al. 2019. A comprehensive approach to analyzing community dynamics using rank abundance curves. - Ecosphere 10: e02881.

Baselga, A. 2010. Partitioning the turnover and nestedness components of beta diversity. - Glob. Ecol. Biogeogr. 19: 134–143.

Bell, T. W. et al. 2020. Three decades of variability in California’s giant kelp forests from the Landsat satellites. - Remote Sens. Environ. 238: 110811.

Brown, B. L. et al. 2016. Compensatory dynamics stabilize aggregate community properties in response to multiple types of perturbations. - Ecology 97: 2021–2033.

Buckley, H. L. et al. 2021. Changes in the analysis of temporal community dynamics data: a 29-year literature review. - PeerJ 9: e11250.

Chalcraft, D. R. 2013. Changes in ecological stability across realistic biodiversity gradients depend on spatial scale. - Glob. Ecol. Biogeogr. 22: 19–28.

Chase, J. M. 2010. Stochastic community assembly causes higher biodiversity in more productive environments. - Science 328: 1388–1391.

Clutton-Brock, T. and Sheldon, B. C. 2010. Individuals and populations: the role of long-term, individual-based studies of animals in ecology and evolutionary biology. - Trends Ecol. Evol. 25: 562–573.

De Cáceres, M. et al. 2019. Trajectory analysis in community ecology. - Ecol. Monogr. 89: e01350.

Diaz, A. et al. 2013. Conservation implications of long-term changes detected in a lowland heath plant metacommunity. - Biol. Conserv. 167: 325–333.

Donohue, I. et al. 2013. On the dimensionality of ecological stability. - Ecol. Lett. 16: 421–429.

Donohue, I. et al. 2016. Navigating the complexity of ecological stability. - Ecol. Lett. 19: 1172–1185.

Dornelas, M. et al. 2014. Assemblage time series reveal biodiversity change but not systematic loss. - Science 344: 296–9.

Edwards, M. et al. 2010. Multi-decadal oceanic ecological datasets and their application in marine policy and management. - Trends Ecol. Evol. 25: 602–610.

Gimona, A. et al. 2012. Woodland networks in a changing climate: Threats from land use change. - Biol. Conserv. 149: 93–102.

Gonzalez, A. and Loreau, M. 2009. The causes and consequences of compensatory dynamics in ecological communities. - Annu. Rev. Ecol. Evol. Syst. 40: 393–414.

Gouhier, T. C. et al. 2010. Synchrony and stability of food webs in metacommunities. - Am. Nat. 175: E16–E34.

Grimm, V. and Wissel, C. 1997. Babel, or the ecological stability discussions: An inventory and analysis of terminology and a guide for avoiding confusion. - Oecologia 109: 323–334.

Harrer, S. L. et al. 2013. Patterns and controls of the dynamics of net primary production by understory macroalgal assemblages in giant kelp forests. - J. Phycol. 49: 248–257.

Hautier, Y. et al. 2018. Local loss and spatial homogenization of plant diversity reduce ecosystem multifunctionality. - Nat. Ecol. Evol. 2: 50–56.

Hillebrand, H. and Kunze, C. 2020. Meta-analysis on pulse disturbances reveals differences in functional and compositional recovery across ecosystems. - Ecol. Lett. 23: 575–585.

Hillebrand, H. et al. 2010. Warming leads to higher species turnover in a coastal ecosystem. - Glob. Change Biol. 16: 1181–1193.

Hillebrand, H. et al. 2018. Decomposing multiple dimensions of stability in global change experiments. - Ecol. Lett. 21: 21–30.

Hughes, B. B. et al. 2017. Long-term studies contribute disproportionately to ecology and policy. - BioScience 67: 271–281.

Ims, R. A. and Steen, H. 1990. Geographical synchrony in microtine population cycles: a theoretical evaluation of the role of nomadic avian predators. - Oikos, 381–387.

Ives, A. R. and Carpenter, S. R. 2007. Stability and diversity of ecosystems. - Science 317: 58–62.

Jost, L. 2007. Partitioning diversity into independent alpha and beta components. - Ecology 88: 2427–2439.

Jost, L. et al. 2011. Compositional similarity and β diversity. - In: Magurran, A. E. and McGill, B. J. (eds), Biological diversity: frontiers in measurement and assessment. Oxford University Press, pp. 66–84.

Lamy, T. et al. 2015. Understanding the spatio-temporal response of coral reef fish communities to natural disturbances: insights from beta-diversity decomposition. - PLOS ONE 10: e0138696.

Lamy, T. et al. 2019. Species insurance trumps spatial insurance in stabilizing biomass of a marine macroalgal metacommunity. - Ecology 100: e02719.

Legendre, P. 2014. Interpreting the replacement and richness difference components of beta diversity. - Glob. Ecol. Biogeogr. 23: 1324–1334.

Legendre, P. 2019. A temporal beta-diversity index to identify sites that have changed in exceptional ways in space–time surveys. - Ecol. Evol. 9: 3500–3514.

Legendre, P. and Gallagher, E. D. 2001. Ecologically meaningful transformations for ordination of species data. - Oecologia 129: 271–280.

Legendre, P. and Legendre, L. 2012. Numerical Ecology. - Elsevier.

Legendre, P. and De Cáceres, M. 2013. Beta diversity as the variance of community data: dissimilarity coefficients and partitioning. - Ecol. Lett. 16: 951–963.

Legendre, P. and Gauthier, O. 2014. Statistical methods for temporal and space-time analysis of community composition data. - Proc. R. Soc. B Biol. Sci. 281: 20132728–20132728.

Leibold, M. A. and Chase, J. M. 2018. Metacommunity ecology. - Princeton University Press.

Leibold, M. A. et al. 2004. The metacommunity concept: a framework for multi-scale community ecology. - Ecol. Lett. 7: 601–613.

Loreau, M. et al. 2003. Biodiversity as spatial insurance in heterogeneous landscapes. - Proc. Natl. Acad. Sci. 100: 12765–12770.

MacArthur, R. 1955. Fluctuations of animal populations and a measure of community stability. - Ecology 36: 533–536.

Magurran, A. E. et al. 2018. Divergent biodiversity change within ecosystems. - Proc. Natl. Acad. Sci. 115: 1843–1847.

Magurran, A. E. et al. 2019. Temporal β diversity—A macroecological perspective. - Glob. Ecol. Biogeogr. 28: 1949–1960.

Margules, C. R. and Pressey, R. L. 2000. Systematic conservation planning. - Nature 405: 243–253.

May, R. M. 1973. Stability and complexity in model ecosystems. - Princeton University Press.

Micheli, F. et al. 1999. The dual nature of community variability. - Oikos 85: 161–169.

Moran, P. A. P. 1953. The statistical analysis of the Canadian Lynx cycle. - Aust. J. Zool. 1: 291–298.

Nguyen, T. H. et al. 2018. A spatial and temporal analysis of forest dynamics using Landsat time-series. - Remote Sens. Environ. 217: 461–475.

Oliver, T. et al. 2010. Heterogeneous landscapes promote population stability. - Ecol. Lett. 13: 473–484.

Orloci, L. 1967. An agglomerative method for classification of plant communities. - J. Ecol. 55: 193–206.

Podani, J. et al. 2013. A general framework for analyzing beta diversity, nestedness and related community-level phenomena based on abundance data. - Ecol. Complex. 15: 52–61.

Post, E. and Forchhammer, M. C. 2004. Spatial synchrony of local populations has increased in association with the recent Northern Hemisphere climate trend. - Proc. Natl. Acad. Sci. 101: 9286–9290.

Rao, C. R. 1995. A review of canonical coordinates and an alternative to correspondence analysis using Hellinger distance. - Qüestiió Quad. Estad. Investig. Oper. 19: 22–63.

Record, S. et al. 2021. Novel insights to be gained from applying metacommunity theory to long-term, spatially replicated biodiversity data. - Front. Ecol. Evol. 8: 612794.

Reed, D. C. 2018. SBC LTER: Reef: Annual time series of biomass for kelp forest species, ongoing since 2000. Environmental Data Initiative. https://doi.org/10.6073/pasta/d5fd133eb2fd5bea885577caaf433b30.

de Roos, A. M. et al. 1998. Pattern formation and the spatial scale of interaction between predators and their prey. - Theoretical Population Biology, 53: 108–130.

Ruhi, A. et al. 2018. Detrimental effects of a novel flow regime on the functional trajectory of an aquatic invertebrate metacommunity. - Glob. Change Biol. 24: 3749–3765.

Shoemaker, L. G. and Melbourne, B. A. 2016. Linking metacommunity paradigms to spatial coexistence mechanisms. - Ecology 97: 2436–2446.

Socolar, J. B. et al. 2016. How should beta-diversity inform biodiversity conservation? - Trends Ecol. Evol. 31: 67–80.

Steiner, C. F. et al. 2013. Population synchrony and stability in environmentally forced metacommunities. - Oikos 122: 1195–1206.

Sturbois, A. et al. 2021. Extending community trajectory analysis: New metrics and representation. - Ecol. Model. 440: 109400.

Tatsumi, S. et al. 2020. Partitioning the colonization and extinction components of beta diversity across disturbance gradients. - Ecology 101: e03183.

Tatsumi, S. et al. 2021. Temporal changes in spatial variation: partitioning the extinction and colonisation components of beta diversity. - Ecol. Lett. 24: 1063–1072.

Tilman, D. 1999. The ecological consequences of changes in biodiversity: a search for general principles. - Ecology 80: 1455–1474.

Tuomisto, H. 2010a. A diversity of beta diversities: Straightening up a concept gone awry. Part 1. Defining beta diversity as a function of alpha and gamma diversity. - Ecography 33: 2–22.

Tuomisto, H. 2010b. A diversity of beta diversities: Straightening up a concept gone awry. Part 2. Quantifying beta diversity and related phenomena. - Ecography 33: 23–35.

Tuomisto, H. et al. 2012. Modelling niche and neutral dynamics: on the ecological interpretation of variation partitioning results. - Ecography 35: 961–971.

Wang, S. and Loreau, M. 2014. Ecosystem stability in space: α, β and γ variability. - Ecol. Lett. 17: 891–901.

Wang, S. and Loreau, M. 2016. Biodiversity and ecosystem stability across scales in metacommunities. - Ecol. Lett. 19: 510–518.

Wang, S. et al. 2017. An invariability-area relationship sheds new light on the spatial scaling of ecological stability. - Nat. Commun. 8: 15211.

Wang, S. et al. 2019. Stability and synchrony across ecological hierarchies in heterogeneous metacommunities: linking theory to data. - Ecography 42: 1200–1211.

Wang, S. et al. 2021. Biotic homogenization destabilizes ecosystem functioning by decreasing spatial asynchrony. - Ecology 102: e03332.

White, L. et al. 2020. Individual species provide multifaceted contributions to the stability of ecosystems. - Nat. Ecol. Evol. 4: 1594–1601.

Whittaker, R. H. 1952. A study of summer foliage insect communities in the Great Smoky Mountains. - Ecol. Monogr. 22: 2–44.

Wilcox, K. R. et al. 2017. Asynchrony among local communities stabilises ecosystem function of metacommunities. - Ecol. Lett. 20: 1534–1545.

Wolda, H. 1981. Similarity indices, sample size and diversity. - Oecologia 50: 296–302.

Xu, Q. et al. 2021. Consistently positive effect of species diversity on ecosystem, but not population, temporal stability. - Ecol. Lett. in press.

Yachi, S. and Loreau, M. 1999. Biodiversity and ecosystem productivity in a fluctuating environment: the insurance hypothesis. - Proc. Natl. Acad. Sci. U. S. A. 96: 1463–1468.

