## Supplementary text 1 for "The dual nature of metacommunity variability" (from another version of this preprint: version 1, 2021-04-11)

**Supplementary text 1. Quantifying temporal compositional variability across spatial scales**

In this supplementary text, we introduce two ways of quantifying temporal compositional variability across spatial scales. By compositional variability we mean changes in species composition that are independent from changes in aggregate community property (e.g. total community biomass). In this framework, we refer to the spatial compositional synchrony component as the key component linking compositional variability across spatial scales, which captures the degree of spatial synchrony in compositional changes across a set of local communities.

**Notation**

We assume that *s* species are surveyed at *m* local communities for *n* time steps. $X_{itj}$ denotes the biomass of species *j*, in local community *i* at time *t*.

**Defining temporal compositional variability of a single community**

The temporal compositional variability (TCV) of a single local community *i* composed of *s* species surveyed *n* times can be computed based on beta diversity following the variance framework developed by Legendre and De Cáceres (2013). In this framework, the TCV of local community *i* represents the total variance in species composition observed over time and is computed as:

${BD}_{i}={SST}/{(n-1)}$	**(1)**

where $SST$is the total sum of squares in species composition, computed as $SST=\sum_{j}^{s} \sum_{t}^{n} {(X_{itj}-\bar{X}_{i.j})}^{2}$, with ${(X_{itj}-\bar{X}_{i.j})}^{2}$ the square of the difference between the biomass of species *j* at time *t*, and the temporal mean biomass of species *j* ($\bar{X}_{i.j}= {\sum_{t}^{n} X_{itj}}/n$). This definition of beta diversity corresponds to the Euclidean distance, which is both inappropriate for beta diversity assessment (Wolda 1981; Legendre and Gallagher 2001), and does not fulfills the “*density invariance*” property (Jost et al. 2011).

Other pairwise dissimilarities than the Euclidean distance can be used to assess beta diversity (Legendre and De Cáceres 2013). In order to properly capture TCV, the computation of beta diversity should be based on a pairwise dissimilarity that fulfill the “*density invariance*” property defined by Jost et al. (2011). Three abundance-based dissimilarity indices fulfil this criterion and correspond to the Whittaker’s index of association (Whittaker 1952), the Hellinger distance (Rao 1995) and the Chord distance (Orlóci 1967). Here, we will show how TCV can be computed based on the Hellinger distance (${BD}^{h}$) and on a time-independent version of the Hellinger distance (time-independent Hellinger distance: ${BD}^{hT}$).

$SST$ corresponding to the Hellinger distance can be obtained directly by computing the same-name transformation on the original data. The Hellinger transformation corresponds to:

$X_{itj}^{h}=\sqrt{{X_{itj}}/{X_{it.}}}$	**(2)**

Where $X_{it.}=\sum_{j}^{s} X_{itj}$ corresponds to the total community biomass observed at time *t* in the local community *i*. In other words, the Hellinger transformation is based on the square root coefficient of species relative frequencies, and thus provides a measure of compositional variability that is independent from differences in total community biomass among observations (“*density invariance*” property [Jost et al. 2011]). TCV based on the Hellinger distance in local community *i* can then be rewritten as:

${BD}_{i}^{h}=\frac{\sum_{j}^{s} \sum_{t}^{n} {(X_{itj}^{h}-\bar{X}_{i.j}^{h})}^{2}}{(n-1)}$	**(3)**

with $\bar{X}_{i.j}^{h}={\sum_{t}^{n} \sqrt{{X_{itj}}/{X_{it.}}}}/n$, the temporal mean of the square root coefficient of the relative frequency of species *j*. ${BD}_{i}^{h}$ can be rewritten as:

${BD}_{i}^{h}=\sum_{j}^{s} v_{ij}^{h}$	**(4)**

With $v_{ij}^{h}=\frac{\sum_{t}^{n} {(X_{itj}^{h}-\bar{X}_{i.j}^{h})}^{2}}{n-1}$ the temporal variance of the Hellinger-transformed biomass of species *j* in local community *i*.

The advantage of beta diversity indices based on the variance framework (Legendre and De Cáceres 2013), such as ${BD}_{i}^{h}$, is that they are comparable among taxonomic groups, as they range between zero, when a local community exhibits the exact same species composition over time, and one, when it exhibits distinct species compositions at each time step. However, while the advantage of using ${BD}^{h}$ to assess TCV in a single community is clear and appealing, it does not behave as one could expect as ${BD}_{i}^{h}$ does not necessarily decrease with increasing spatial scale. For instance, let’s assume that we want to assess the TCV of two local communities across spatial scales, and that the two local communities are made of two distinct species fluctuating over time (Table 1). In this example, ${BD}_{1}^{h}$ and ${BD}_{2}^{h}$of in communities 1 and 2 respectively will both be null, but it is > 0 at larger spatial scales (*i.e.* for the combination of the two communities corresponding to the gamma scale, ${BD}_{\gamma}^{h}$, see below). It is preferable for TCV to decrease with increasing spatial scale (*i.e.*, alpha $\geq$ gamma), such as in the case of aggregate variability (Wang and Loreau 2014), or species richness (Scheiner 2003). In this case, ${BD}_{i}^{h}$ does not always decrease with increasing spatial scale because the Hellinger transformation standardized each observation in a time dependent way (*i.e.*, dividing by $X_{it.}$).

Therefore, we introduce a modified Hellinger transformation to obtain a time-independent normalization of the original data as:

$X_{itj}^{hT}=\sqrt{{X_{itj}}/{X_{i..}}}$	**(5)**

Where $X_{i..}$ corresponds to the total community biomass observed across all time steps in local community *i* ($X_{i..}=\sum_{j}^{s} \sum_{t}^{n} X_{itj}$). Instead of normalizing the biomass of each species by the total community biomass observed at each time step, we normalize species biomass by the total community biomass observed across all time steps. Hence it corresponds to the square root coefficient of species relative frequencies with respect to the biomass of the whole local community. TCV based on the time-independent Hellinger transformation is:

${BD}_{i}^{hT}=\frac{\sum_{j}^{s} \sum_{t}^{n} \left( X_{itj}^{hT}-\bar{X}_{i.j}^{hT} \right)^{2}}{\left( n-1 \right)}$	**(6)**

with $\bar{X}_{i.j}^{hT}={\sum_{t}^{n} \sqrt{{X_{itj}}/{X_{i..}}}}/n$, the temporal mean of the square root coefficient of the time-independent relative frequency of species *j*. ${BD}_{i}^{hT}$ can be rewritten as:

${BD}_{i}^{hT}=\frac{1}{X_{i..}}\frac{\sum_{j}^{s} \sum_{t}^{n} \left( \sqrt{X_{itj}}-\frac{\sum_{t}^{n} \sqrt{X_{itj}}}{n} \right)^{2}}{\left( n-1 \right)}=\frac{\sum_{j}^{s} v_{ij}^{hT}}{X_{i..}}$	**(7)**

With $v_{ij}^{hT}=\frac{\sum_{t}^{n} \left( \sqrt{X_{itj}}-\bar{\sqrt{X_{i.j}}} \right)^{2}}{n-1}$ the temporal variance of the square root coefficient biomass of species *j* in local community *i*.

In our previous example of two communities composed of two distinct species (Table 1), ${BD}_{1}^{hT}$ and ${BD}_{2}^{hT}$are greater than zero for both local communities as their modified species frequencies are now different from one. More generally, ${BD}_{i}^{hT}$ decreases with increasing spatial scales (*i.e.*, alpha $\geq$ gamma). Therefore, this new definition might be more operational to study TCV across spatial scales as compared to ${BD}_{i}^{h}$. However, ${BD}_{i}^{hT}$ can be greater than one. Hence both definition provide interesting insights into temporal compositional changes, one rooted into the variance framework of beta diversity, the other is more suited to deal with multiple scales. In addition, ${BD}_{i}^{h}$ can be used when only species relative frequency is available. We further discuss these two indices below.

**Local scale temporal compositional variability**

We define local scale of TCV as the weighted average of TCV across the *m* local communities. Local scale TCV based on the Hellinger distance is:

${BD}_{\alpha}^{h}=\sum_{i}^{m} \frac{X_{i..}}{X_{...}}{BD}_{i}^{h}=\sum_{i}^{m} \frac{X_{i..}}{X_{...}}\sum_{j}^{s} v_{ij}^{h}$	**(8)**

And local scale TCV based on the time-independent Hellinger transformation is:

${BD}_{\alpha}^{hT}=\sum_{i}^{m} \frac{X_{i..}}{X_{...}}{BD}_{i}^{hT}=\frac{\sum_{i}^{m} \sum_{j}^{s} v_{ij}^{hT}}{X_{...}}$	**(9)**

With $\frac{X_{i..}}{X_{...}}$ the weight of local community *i*. Local communities are weighted according to their contributions to the total metacommunity biomass with $X_{...}=\sum_{i}^{m} \sum_{t}^{n} \sum_{j}^{s} X_{itj}$ and $X_{i..}=\sum_{t}^{n} \sum_{j}^{s} X_{itj}.$

**Regional scale compositional variability**

We define the regional scale of temporal compositional variability based on the Hellinger distance as:

${BD}_{\gamma}^{h}=\frac{{\sum_{j}^{s} \sum_{t}^{n} (X_{.tj}^{h}-\bar{X}_{..j}^{h})}^{2}}{(n-1)}=\sum_{j}^{s} v_{Tj}^{h}$	**(10)**

with $X_{.tj}^{h}=\sqrt{{\sum_{i}^{m} X_{itj}}/{X_{.t.}}}$ the square root coefficient of the regional frequency of species *j* at time *t*, and $\bar{X}_{..j}^{h}=\frac{\sum_{t}^{n} X_{.tj}^{h}}{n}$ the temporal mean of the square root coefficient of the regional frequencies of species *j*. $v_{Tj}^{h}=\frac{\sum_{t}^{n} {(X_{.tj}^{h}-\bar{X}_{..j}^{h})}^{2}}{n-1}$ corresponds to the temporal variance of the Hellinger-transformed regional biomass of species *j*.

We define the regional scale of temporal compositional variability based on the time-independent Hellinger transformation:

${BD}_{\gamma}^{hT}=\frac{{\sum_{j}^{s} \sum_{t}^{n} (X_{.tj}^{hT}-\bar{X}_{..j}^{hT})}^{2}}{(n-1)}=\frac{\sum_{j}^{s} v_{Tj}^{hT}}{X_{...}}$	**(11)**

with $X_{.tj}^{hT}=\sqrt{{\sum_{i}^{m} X_{itj}}/{X_{...}}}$ the square root coefficient of the time-independent regional frequency of species *j*, and $\bar{X}_{..j}^{hT}=\frac{\sum_{t}^{n} X_{.tj}^{hT}}{n}$ the temporal mean of the square root coefficient of the time-independent regional frequencies of species *j*. $v_{Tj}^{hT}=\frac{\sum_{t}^{n} \left( \sqrt{X_{.tj}}-\bar{\sqrt{X_{..j}}} \right)^{2}}{n-1}$ corresponds to the temporal variance of the square root coefficient of the regional biomass of the *j*^th^ species.

**Multiplicative partitioning of temporal compositional variability across spatial scales**

TCV at the regional scale (${BD}_{\gamma}^{h}$ and ${BD}_{\gamma}^{hT}$) can be partitioned multiplicatively into a local scale (${BD}_{\alpha}^{h}$ and ${BD}_{\alpha}^{hT}$) and a spatial component $({BD}_{\varphi}^{h}$ and ${BD}_{\varphi}^{hT}$) as:

${BD}_{\gamma}^{h}={BD}_{\alpha}^{h}.{BD}_{\varphi}^{h}$	**(12)**

And

${BD}_{\gamma}^{hT}={BD}_{\alpha}^{hT}.{BD}_{\varphi}^{hT}$	**(13)**

${BD}_{\varphi}^{h}$ and ${BD}_{\varphi}^{hT}$ are the spatial component scaling compositional variability across spatial scales. An important distinction between the two approaches is that ${BD}_{\varphi}^{h}$ does not entirely correspond to spatial synchrony index that ranges from zero to one, as it is defined only as the ratio between ${BD}_{\gamma}^{h}$ and ${BD}_{\alpha}^{h}$. Indeed, since ${BD}_{\gamma}^{h}$ can be smaller than ${BD}_{\alpha}^{h}$, ${BD}_{\varphi}^{h}$ can be greater than one. ${BD}_{\varphi}^{h}$ defined as the ratio between gamma and alpha compositional variability can be rewrite as:

${BD}_{\varphi}^{h}=\frac{{BD}_{\gamma}^{h}}{{BD}_{\alpha}^{h}}= \frac{\sum_{j}^{s} v_{Tj}^{h}}{\sum_{i}^{m} \frac{X_{i..}}{X}\sum_{j}^{s} v_{ij}^{h}}$	**(14)**

On the contrary, ${BD}_{\varphi}^{hT}$ is more appropriate when dealing with multiple spatial scales as it will yields a synchrony metric ranging from zero. Indeed ${BD}_{\varphi}^{hT}$ can be rewritten as:

${BD}_{\varphi}^{hT}=\frac{{BD}_{\gamma}^{hT}}{{BD}_{\alpha}^{hT}}= \frac{\sum_{j}^{s} v_{Tj}^{hT}}{\sum_{i}^{m} \sum_{j}^{s} v_{ij}^{hT}}$	**(15)**

If we define the spatial synchrony of species *j* as:

$\varphi_{(j)}^{hT}=\frac{v_{Tj}^{hT}}{\sum_{i}^{m} v_{ij}^{hT}}$	**(16)**

then:

${BD}_{\varphi}^{hT}= \frac{\sum_{j}^{s} \sum_{i}^{m} v_{ij}^{hT}.\varphi_{(j)}^{hT}}{\sum_{i}^{m} \sum_{j}^{s} v_{ij}^{hT}}=\sum_{j}^{s} \frac{\sum_{i}^{m} v_{ij}^{hT}}{\sum_{i}^{m} \sum_{j}^{s} v_{ij}^{hT}}.\varphi_{(j)}^{hT}=\sum_{j}^{s} w_{j}.\varphi_{(j)}^{hT}$	**(17)**

with $w_{j}= {\sum_{i}^{m} v_{ij}^{hT}}/{\sum_{j}^{s} \sum_{i}^{m} v_{ij}^{hT}}$ reflecting species weight. Hence, ${BD}_{\varphi}^{hT}$ corresponds to the weighted average of species-level spatial synchrony. $\varphi_{(j)}^{hT}$, however, differ from the classic form of other synchrony indices, which usually take the form ${v_{X}}/{\left( \sum\sqrt{v_{x}} \right)^{2}}$ (Loreau and de Mazancourt 2008), such as in the case of the spatial synchrony index of the aggregate community metric (Wang and Loreau 2014). A notable difference is that species biomasses are square root transformed before computing their temporal variances. The denominator of $\varphi_{(j)}^{hT}$, $\sum_{i=1}^{m} v_{ij}^{hT}$, corresponds to the sum across all local populations of the temporal variances of square root coefficient biomass of species *j*, and the numerator, $v_{Tj}^{hT}$ corresponds to the temporal variance of the square root coefficient of its summed biomass across all local communities. ${BD}_{\varphi}^{hT}$ ranges between min($\varphi_{(j)}^{hT}$) and max($\varphi_{(j)}^{hT}$).

**Table SM1**. Comparison between ${BD}^{h}$ and ${BD}^{hT}$. Simple case study of two communities that do not share any species in common.

|  | **Community 1** | |  | **Community 2** | |
| --- | --- | --- | --- | --- | --- |
| **Time** | **Species 1** | **Species 2** |  | **Species 1** | **Species 2** |
| **1** | 2 | 0 |  | 0 | 8 |
| **2** | 4 | 0 |  | 0 | 4 |
| **3** | 6 | 0 |  | 0 | 2 |
| **4** | 3 | 0 |  | 0 | 5 |
| **5** | 5 | 0 |  | 0 | 2 |
| **6** | 1 | 0 |  | 0 | 5 |
| *Hellinger distance* | | | | |  |
| ${BD}_{1}^{h}$ | 0 |  |  | ${BD}_{2}^{h}$ | 0 |
| ${BD}_{\alpha}^{h}$ | 0 |  |  |  |  |
| ${BD}_{\gamma}^{h}$ | 0.069 |  |  |  |  |
| ${BD}_{\varphi}^{h}$ | Infinite |  |  |  |  |
| *Time-independent Hellinger transformation* | | | | |  |
| ${BD}_{1}^{hT}$ | 0.014 |  |  | ${BD}_{2}^{hT}$ | 0.011 |
| ${BD}_{\alpha}^{hT}$ | 0.013 |  |  |  |  |
| ${BD}_{\gamma}^{hT}$ | 0.012 |  |  |  |  |
| ${BD}_{\varphi}^{hT}$ | 0.990 |  |  |  |  |

Table SM2. Notations summary for TCV bases on the Hellinger distance. See Table 2 of the main text for the time-independent Hellinger transformation.

| **Symbol** | **Eq.** | **Description** |
| --- | --- | --- |
| $X_{itj}$ |  | Biomass, or abundance, of species *j* in community *i* at time *t* |
| $X_{it.}=\sum_{j}^{s} X_{itj}$ |  | Total aggregate metric of community *i* at time *t* |
| $X_{itj}^{h}=\sqrt{{X_{itj}}/{X_{it.}}}$ | (2) | Hellinger transformation of the biomass, or abundance, of species *j* in community *i* at time *t*, corresponding to the square root coefficient of species relative |
| $X_{i..}=\sum_{t}^{n} \sum_{j}^{s} X_{itj}$ | (3) | Total aggregate metric (*i.e.* total biomass) of community *i* |
| **Local scale compositional variability based on the Hellinger distance** | | |
| $v_{ij}^{h}=\frac{\sum_{t}^{n} {(\sqrt{{X_{itj}}/{X_{it.}}}-\bar{\sqrt{{X_{itj}}/{X_{it.}}}})}^{2}}{n-1}$ |  | Temporal variance of the Hellinger-transformed biomass of species *j* in community *i* |
| ${BD}_{i}^{h}=\sum_{j}^{s} v_{ij}^{h}$ | (4) | Temporal compositional variability of community *i*, corresponding to the beta diversity based on the Hellinger distance of community *i* |
| ${BD}_{\alpha}^{h}=\sum_{i}^{m} \frac{X_{i..}}{X_{...}}\sum_{j}^{s} v_{ij}^{h}$ | (8) | Local scale aggregate variability, defined as the weighted average of local compositional changes based on the Hellinger distance |
| **Regional scale compositional variability based on the Hellinger distance** | | |
| $X_{.tj}=\sum_{i}^{m} X_{itj}$ |  | Total aggregate metric of species *j* at time *t* in the whole metacommunity |
| $X_{.t.}=\sum_{i}^{m} \sum_{j}^{s} X_{itj}$ |  | Total aggregate metric at time *t* in the whole metacommunity |
| $X_{...}=\sum_{i}^{m} \sum_{t}^{n} \sum_{j}^{s} X_{itj}$ |  | Total aggregate metric of the whole metacommunity |
| $v_{Tj}^{h}=\frac{\sum_{t}^{n} {(\sqrt{{X_{.tj}}/{X_{.t.}}}-\bar{\sqrt{{X_{.tj}}/{X_{.t.}}}})}^{2}}{n-1}$ |  | Temporal variance of the square root coefficient of the regional biomass of species *j* |
| ${BD}_{\gamma}^{h}=\sum_{j}^{s} v_{Tj}^{h}$ | (19) | Regional scale compositional variability, or compositional variability of the whole metacommunity, corresponding to the beta diversity based on the Hellinger distance of the whole metacommunity |
| **Spatial ratio based on the Hellinger distance** | | |
| ${BD}_{\varphi}^{h}=\frac{\sum_{j}^{s} v_{Tj}^{h}}{\sum_{i}^{m} \frac{X_{i..}}{X}\sum_{j}^{s} v_{ij}^{h}}$ | (23) | Spatial component scaling compositional variability between ${BD}_{\alpha}^{h}$ and ${BD}_{\gamma}^{T}$. Does not correspond to the synchrony definition of Loreau and de Mazancourt (2008) |

**References**

Jost L, Chao A, Chazdon RL (2011) Compositional similarity and β (beta) diversity. In: Biological diversity: frontiers in measurement and assessment, A. Magurran y BJ McGill (eds.). Oxford University Press, New York. pp 66–87

Legendre P, De Cáceres M (2013) Beta diversity as the variance of community data: dissimilarity coefficients and partitioning. Ecol Lett 16:951–963. doi: 10.1111/ele.12141

Legendre P, Gallagher ED (2001) Ecologically meaningful transformations for ordination of species data. Oecologia 129:271–280. doi: 10.1007/s004420100716

Orlóci L (1967) An agglomerative method for classification of plant communities. J Ecol 50:193–205. doi: 10.2307/2257725

Rao CR (1995) A review of canonical coordinates and an alternative to correspondence analysis using Hellinger distance.

Scheiner SM (2003) Six types of species-area curves. Glob Ecol Biogeogr 12:441–447. doi: 10.1046/j.1466-822X.2003.00061.x

Wang S, Loreau M (2014) Ecosystem stability in space: α, β and γ variability. Ecol Lett. doi: 10.1111/ele.12292

Whittaker RH (1952) A study of summer foliage insect communities in the Great Smoky Mountains. Ecol Monogr 22:1–44.

Wolda H (1981) Similarity indices, sample size and diversity. Oecologia 50:296–302. doi: 10.1007/BF00344966
